# PML nuclear bodies orchestrate the storage and degradation of aggregated HBc in the nucleus and reduce CAM-A-induced apoptosis

**DOI:** 10.64898/2026.06.29.735234

**Authors:** Vaclav Janovec, Laura Meiss-Heydmann, Valerio Taverniti, Christos Satratzemis, Jan Weber, Barbora Lubyová, Ivan Hirsch, Joachim Lupberger, Hannah Vanrusselt, Yannick Debing, Thomas F. Baumert, Eloi R. Verrier

## Abstract

The lack of effective anti-hepatitis B virus (HBV) therapies highlights the need for a new type of treatment that targets different stages of the viral life cycle. The HBV core protein (HBc) is a critical component of this cycle. Various capsid assembly modulators (CAMs) have been developed to target the HBc and inhibit HBV replication. We recently described a subset of capsid assembly modulators (CAMs) that induce the formation of aberrant structures from the HBc in the nucleus, leading to cell death via annexin A1 (ANXA1)-driven apoptosis. Thus, we further elucidated the mechanism of HBc aggregation in the nucleus, with a particular focus on the interplay between nuclear HBc aggregates and PML nuclear bodies. We found that long-term treatment with CAM-A induced the formation of enlarged PML bodies, approximately 1–2 µm in diameter, that accumulated aggregated HBc. PML silencing in HBc-overexpressing HepG2-NTCP cells led to a dramatic increase in apoptosis following CAM-A-induced HBc aggregation, which was associated with elevated ANXA1. Next, we showed that PML nuclear bodies orchestrate proteasomal degradation of nuclear HBc aggregates via sumoylation-dependent recruitment of RNF4. Collectively, our results suggest that PML nuclear bodies act as storage compartments for aggregated HBc proteins in the nucleus, thereby counteracting the apoptotic elimination of cells. Further study of PML function and the targeting of PML nuclear bodies in HBV-infected hepatocytes could reveal new ways to enhance the effectiveness of CAMs.

## Introduction

Despite the availability of a hepatitis B virus (HBV) vaccine, nearly 300 million people still suffer from chronic HBV infection (Glebe et al., 2021). Chronic HBV is a significant global health concern due to its potential to lead to hepatocellular carcinoma or liver failure. Treatment with nucleoside analogues and interferon-alpha has shown a variable degree of effectiveness, with only a small percentage of patients achieving a cure (Kennedy et al., 2025). Therefore, a novel type of treatment that targets various steps in the HBV life cycle is necessary to achieve sustainable control of HBV.

HBV belongs to the *Hepadnaviridae* family of enveloped viruses, which have partially double-stranded DNA that is replicated through reverse transcription. HBV encodes several proteins, one of which is the core antigen (HBc), that play an important role in various steps of the viral life cycle (Niklasch et al., 2021). Most importantly, HBc is necessary for the encapsidation of the viral pregenomic RNA (pgRNA), which serves as a template for reverse transcription when it is encapsidated by HBc together with the viral polymerase (Lott et al., 2000). The phosphorylation and dephosphorylation of the C-terminal domain of HBc play a critical role in the regulation of pgRNA encapsidation and reverse transcription (Zhao et al., 2018; Hsieh et al., 2025). Moreover, HBc phosphorylation plays a role in subsequent disassembly during viral entry (Luo et al., 2020). Therefore, HBc represents a promising target for viral therapy.

Various compound targeting HBV capsid assembly were developed and clinically evaluated. Capsid assembly modulators (CAMs) affect the dynamics of capsid assembly, leading to the formation of abnormal structures or empty capsids. Two main categories are described based on phenotype: CAM-A inducing assembly of aberrant capsids with different morphologies and CAM-E forming empty capsids resembling wild-type capsids but lacking HBV genome (Taverniti et al., 2022). We recently described a subset of CAM-A that causes the nuclear accumulation of HBc aggregates, leading to cell death via ANXA1-driven apoptosis (Kum et al., 2023; Taverniti et al., 2024). CAM-induced apoptosis depends on *de novo* HBc expression and the amount of HBc (Berke et al., 2024). PML nuclear bodies represent one of the cellular protective systems against nuclear aggregated proteins (Gärtner and Muller, 2014). PML orchestrate degradation via RNF4-mediated ubiquitination of protein aggregates followed by proteasome degradation (Guo et al., 2014). Besides, PML nuclear bodies also play a role in apoptosis (Wang et al., 1998). Despite several indications that CAM-A-induced HBc aggregates associate with PML nuclear bodies the role of this association and connection with apoptosis is currently unclear (Vanrusselt et al., 2023; Huber et al., 2018).

In this study, we explored the intricate relationship between PML nuclear bodies and HBc aggregates within the nucleus, with the aim of further elucidating the mechanism of CAM-mediated apoptosis in HBc-containing hepatocytes.

## Results

### CAM-A induce nuclear aggregates that rapidly associate with PML nuclear bodies

To assess whether various CAMs induce nuclear aggregates and their association with PML nuclear bodies, we used a HepG2 cell line, that stably expresses HBc-HA. HA-tag staining allows us to very specifically detect the aggregation effect of CAM-A by confocal microscopy, where a widely distributed signal representing HBc dimers and capsids rapidly accumulates in specific foci after CAM-A treatment, representing aggregated HBc. Thus, confirming that CAMs shift the balance between free core dimer to fully assembled empty capsids or aberrant HBc structures (Berke et al., 2017). We found that various CAM-A treatments resulted in the association of PML nuclear bodies with aggregated HBc (Figure 1A). Moreover, we confirmed that nuclear HBc aggregation is specific to CAM-A and not to CAM-E (supplementary Figure 1), as we previously described (Taverniti et al., 2024). It has previously been published that CAM-A also induces initial cytoplasmic accumulation of HBc, which is effectively cleared by p62-mediated macroautophagy (Lin et al., 2022). Thus, we analysed the kinetics of HBc aggregation in HBc-expressing cells. We confirmed that, eight hours after CAM-A treatment, the majority of HBc aggregates were present in the cytoplasm, although nuclear accumulation of HBc aggregates and association with PML was also observed (Figure 1B). After 16 hours, most of the aggregates were localised in the nucleus whereas most cytoplasmic aggregates were degraded. Therefore, we hypothesise that CAM-A induces rapid HBc aggregation that is efficiently cleared from the cytoplasm, but not from the nucleus.

**Figure 1.**
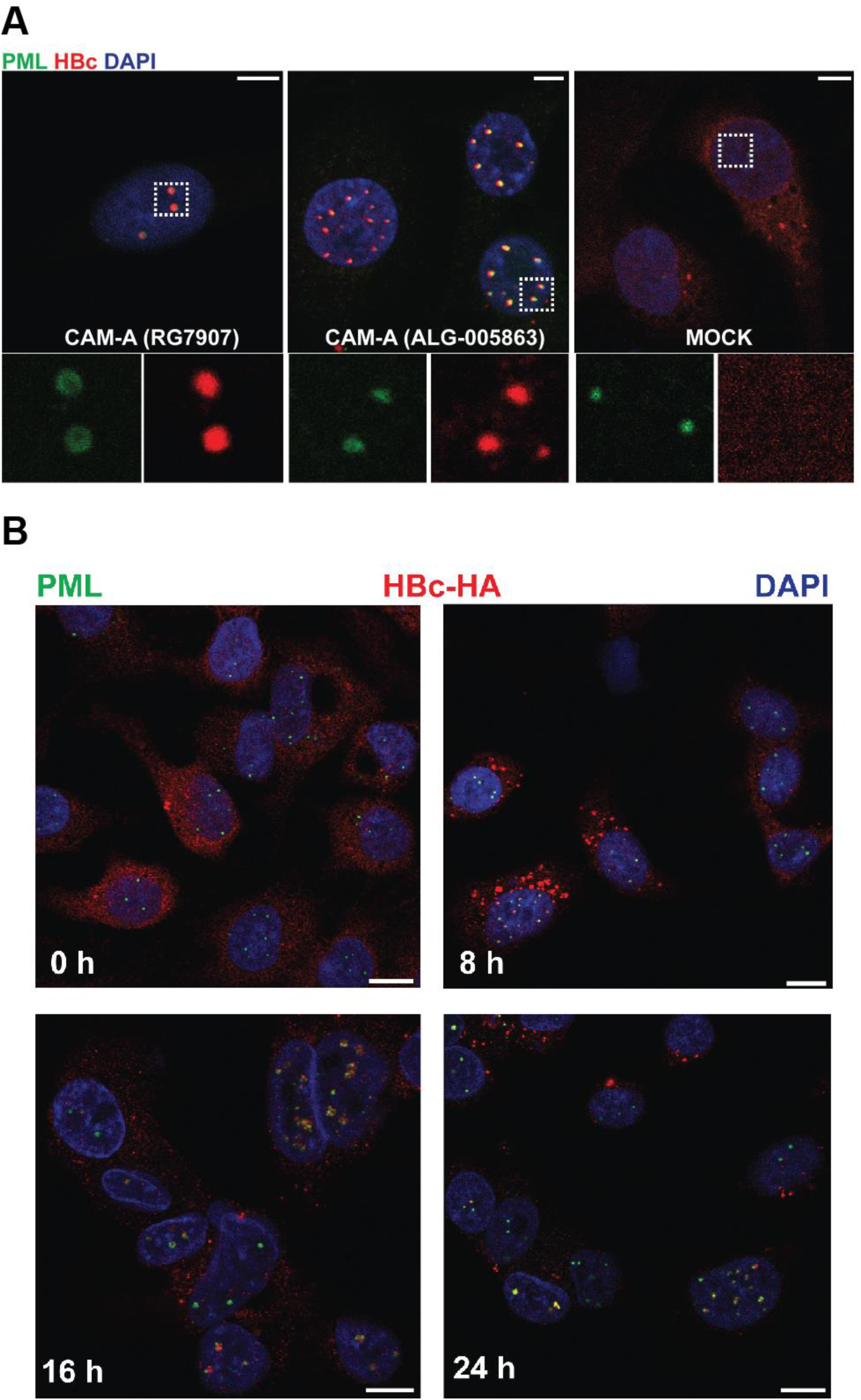
CAM-A mediated HBc aggregation in HepG2-HA-HBc cells. (A) HepG2-HBc-HA cells were treated with CAM-A (RG7907, 1 µM; 24 hours) or MOCK (DMSO) and stained with antibodies against PML (green) and HA-Tag (red). Nuclei were counterstained with DAPI (blue). White squares show enlarged regions of PML and HA-HBc colocalizations analysed by confocal microscopy. Bar = 5 μm (B) HepG2-HA-HBc cells were treated with RG7907 (1 µM) and fixed at various times after the treatment (0, 8, 16 and 24 hours) as indicated and analysed by confocal microscopy. PML (green) and HA-Tag (red) were stained with specific antibodies and DNA by DAPI (blue). Bars = 10 μm.

### Long-term CAM-A treatment leads to the accumulation of aggregated HBc in large ring-shaped PML nuclear bodies in HBV infected HepG2-NTCP cells

To confirm the data obtained in our HBc-overexpression system and avoid the potential influence of the HA-tag on the targeting of HBc aggregates to PML nuclear bodies, we tested the association of HBc aggregates with PML nuclear bodies in HBV-infected cells. For this, we used HepG2-NTCP cells infected with HBV and a capsid-specific antibody that binds to fully established capsids and can detect aggregated HBc (Lubyova et al., 2025). As CAM-A may also affect the early stages of HBV infection (Berke et al., 2017), HepG2-NTCP cells were first infected with HBV, and CAM-A treatment was initiated three days post-infection. Infected cells were analysed 6 days after the start of CAM-A treatment. Our results showed that CAM-A treatment induced formation of HBc nuclear aggregates in HBV-infected HepG2-NTCP cells that associated with PML nuclear bodies (Figure 2A). No association was observed between HBV capsids and PML nuclear bodies in mock-treated HBV-infected HepG2-NTCP cells (supplementary Figure 2). Thus, we confirmed the association between CAM-A-induced HBc aggregates and PML nuclear bodies in HBV-infected cells (Huber et al., 2018). Further analysis of the PML body structure revealed that CAM-A treatment induced formation of large ring-shaped PML bodies with a size of 1-2 µm that enclosed aggregated core (Figure 2B). To further characterize the formation of enlarged PML nuclear bodies, we performed Z-stack imaging of HBc and PML using confocal microscopy. Z-stack imaging confirmed that various numbers of enlarged PML bodies with size 1-2 µm were formed in the nucleus of HBV-infected cells (Figure 2C and supplementary video S1), potentially serving as a nuclear storage compartment for aggregated HBc.

**Figure 2.**
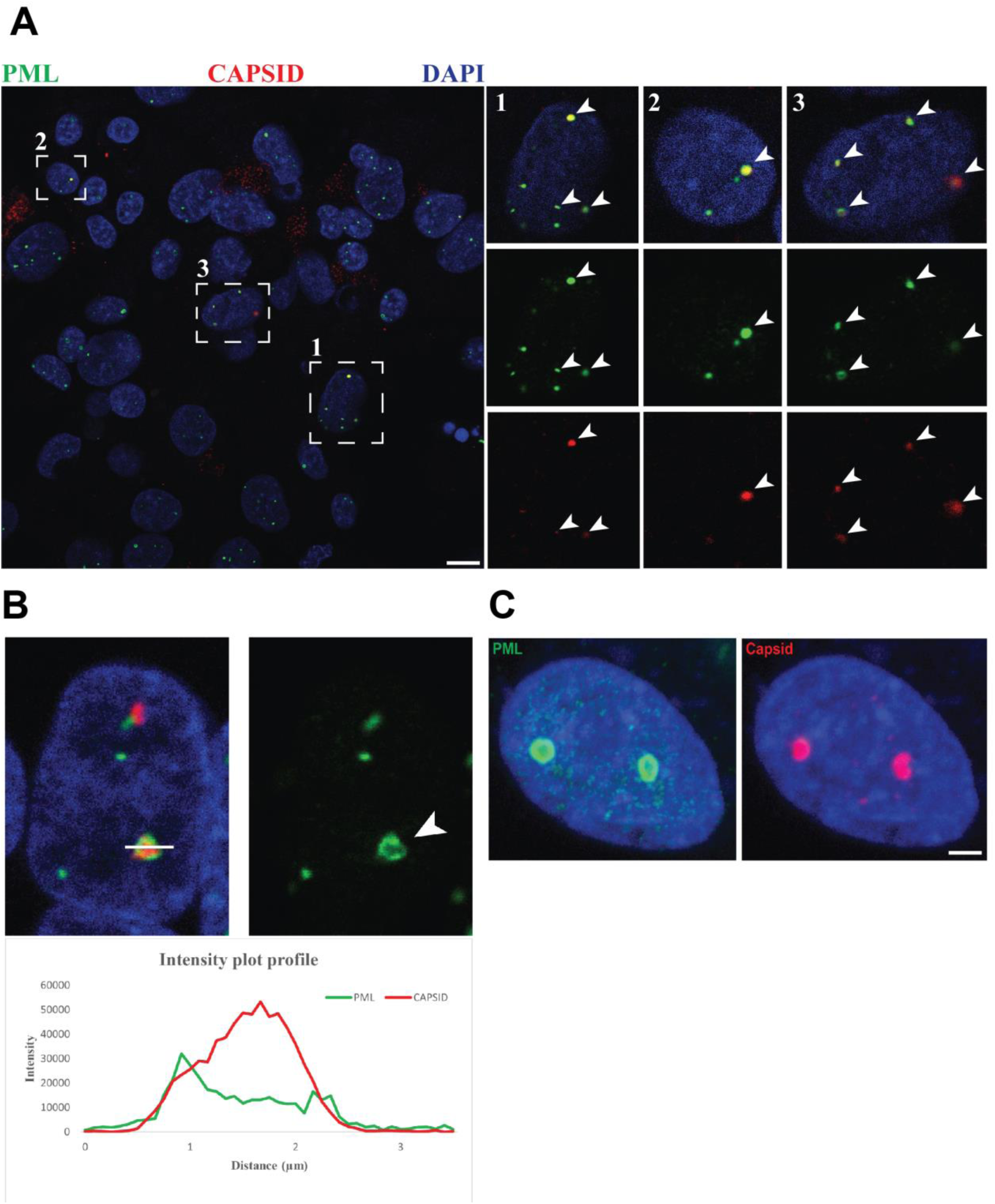
Formation of ring-shaped PML bodies containing aggregated HBc in HBV-infected HepG2-NTCP cells. (A) HepG2-NTCP cells were infected with HBV (1000GE/cell) and treated with CAM-A (ALG-005863; 1 µM; 6 days) and stained with antibodies against PML (green) and HBV capsid (red). Nuclei were counterstained with DAPI (blue). White squares and white arrows show enlarged regions of PML and HBV capsid colocalizations analyzed by confocal microscopy. Bar = 10 μm. (B) Intensity profile analysis of a PML nuclear body containing HBc aggregate by confocal microscopy. The white line indicates the area of interest for intensity profile analysis (shown below), and the white arrow indicate the structure of enlarged PML (green) in single channel image. DNA was stained by DAPI (blue). (C) Z-stack confocal sections of PML nuclear bodies positive for HBc aggregates in an individual cell. Maximum intensity projection was used to present 3D data into 2D image. PML (green) and HBV capsid (red) were stained with specific antibodies and DNA by DAPI (blue). Bar = 2 μm

### PML silencing increases apoptosis in HBc-expressing cells

We previously demonstrated that CAM-A treatment leads to HBc-aggregation-dependent hepatocyte death (Taverniti et al., 2024; Kum et al., 2023). To find out more about the link between CAM-A-induced aggregation in the nucleus and PML nuclear bodies, we studied how PML silencing affects apoptosis (Caspase-3/7 activity) in HBc-expressing cells (Figure 3A). PML protein exists in two states in the nucleus, as free, diffused PML proteins in the nucleoplasm and as components of nuclear bodies (Dorosz et al., 2025). Thus, we first verified by confocal microscopy that PML silencing (supplementary Figure 3) leads to disruption of assembled PML nuclear bodies (Figure 3B). Then, we used HBc-expressing cells and measured Caspase 3/7 activity on day 3 post CAM-A treatment when apoptosis reaches its peak, as we previously described (Taverniti et al. 2024). Our results showed that PML silencing led to an increase in apoptosis in CAM-A treated cells that express wild type HBc, but not in cells expressing the HBc T33N mutant that is resistant to CAM-A treatment (Figure 3C). Our results indicate that PML bodies protects against CAM-A-induced cell death in HBc expressing cells.

**Figure 3.**
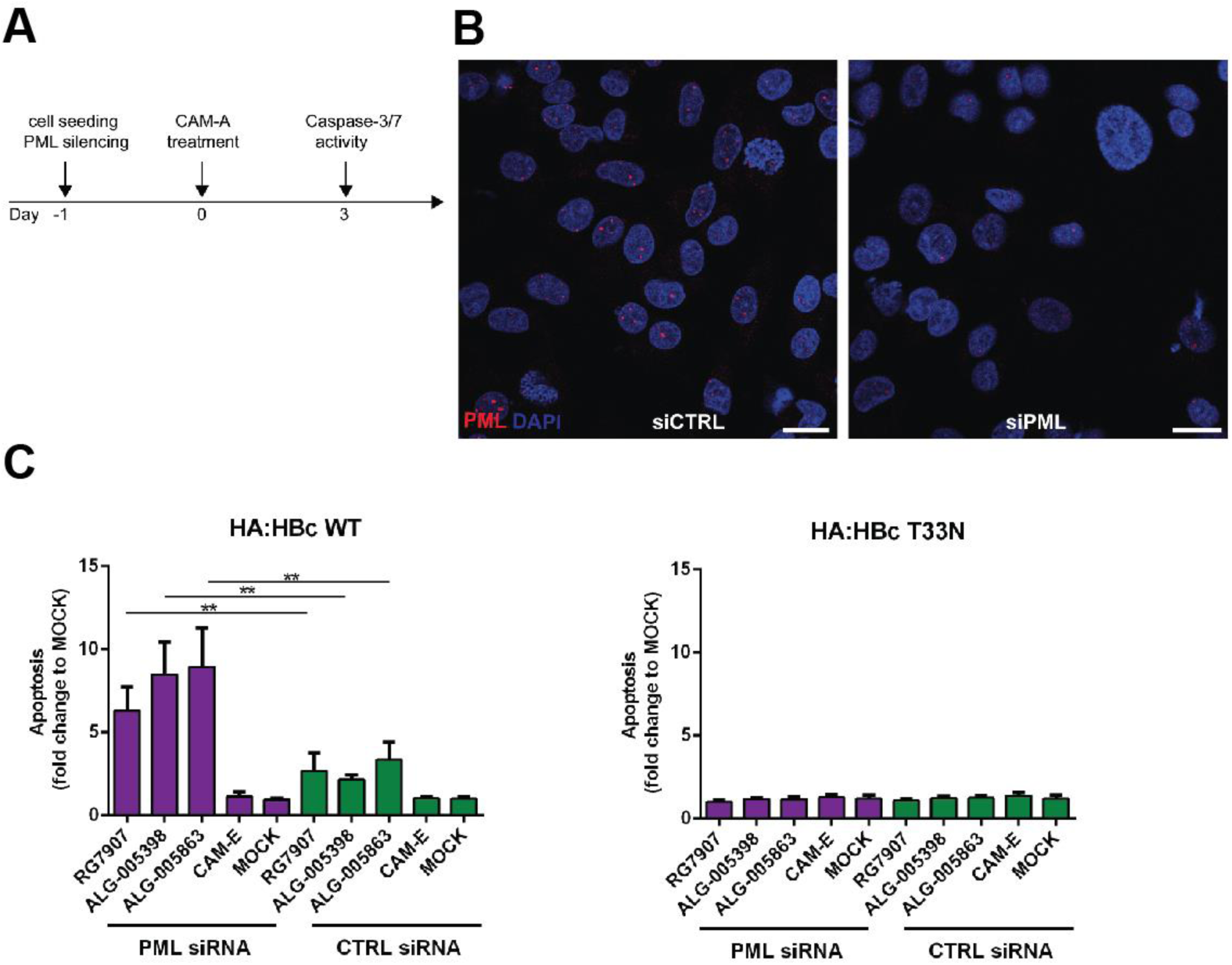
PML silencing leads to an increase of apoptosis in CAM-A-treated HBc-expressing cells. (A) HepG2-NTCP expressing HBc-HA WT or HBc-HA T33N mutant were transfected with siRNA targeting PML or non-targeting siRNA. Then the cells were treated with various CAM-A as indicated. Compound B (Aligos Therapeutics) served as a CAM-E control. (B) The verification of PML silencing by confocal microscopy. PML was stained by specific antibody (red) and nucleus by DAPI (blue). Bars = 10 μm. (C) The effect of PML silencing on the apoptosis was determined by a caspase 3/7 reporter assay. Data are expressed as means of two independent experiments. Levels of significance: ∗∗p <0.01(two-tailed Mann–Whitney U test).

### PML silencing leads to an increase in HBc aggregate size and ANXA-1 expression

Next, we asked whether the PML bodies disruption might affect the CAM-A-mediated aggregation of HBc. To this end, we analysed the shape of nuclear HBc aggregates in CAM-A treated HepG2-NTCP cells expressing HBc after PML knockdown. Surprisingly, we observed, that PML silencing leads to the accumulation of the larger HBc-aggregates compared to the control siRNA (Figure 4A-B). We previously published that CAM-A-induced HBc nuclear aggregation led to the upregulation of ANXA1 expression, and that silencing of ANXA1 expression delayed cell death and apoptosis (Taverniti et al., 2024). Therefore, we analysed the effect of PML silencing on ANXA1 expression in CAM-A-treated HepG2-NTCP expressing HBc. PML silencing increased the number of ANXA1-positive cells after CAM-A treatment, further confirming the protective role of PML bodies against CAM-A-induced HBc aggregation and apoptosis. (Figure 4C).

**Figure 4.**
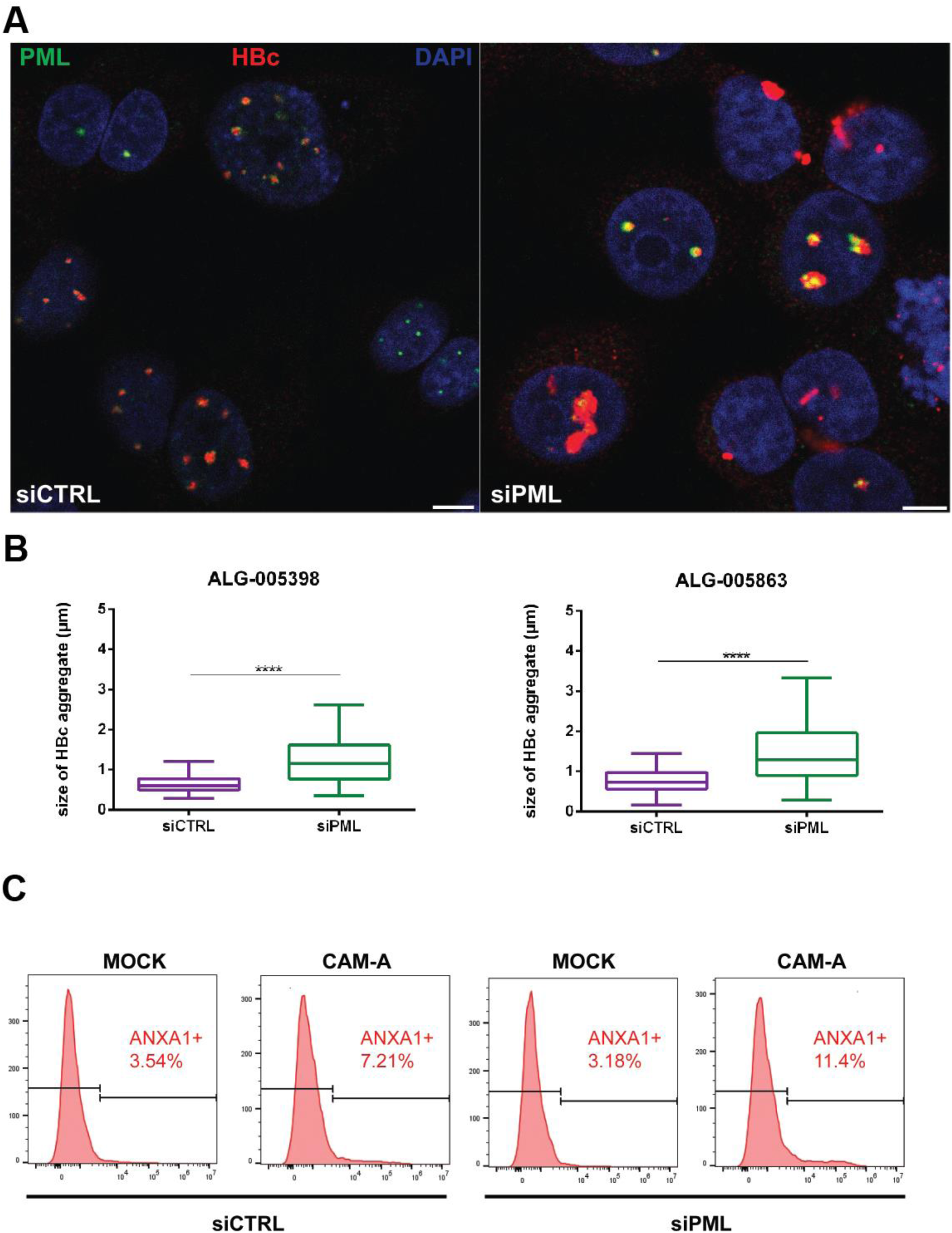
Disruption of PML bodies leads to an increase in HBc nuclear aggregate size and ANXA-1 expression. (A) HepG2-NTCP expressing HBc-HA were transfected with siRNA targeting PML expression or non-targeting siRNA and then treated with CAM-A (ALG-005398, 1 µM, 24 hours). PML (green) and HBc (red) were stained by specific antibodies and nucleus by DAPI (blue) and visualized using confocal microscopy. Bars = 5 μm. (B) The size of HBc aggregates (ALG-005398 or ALG-005863, 1 µM, 24 hours) was evaluated from confocal images using Fiji. The data are expressed as the mean size of aggregated HBc from individual cells. Levels of significance: ∗∗∗∗p <0.0001(two-tailed Mann–Whitney U test). (B) ANXA1 expression in CAM-A treated HepG2-HBc-HA cells (ALG-005398, 1 µM, 3 days) was assessed by flow cytometry using a specific ANXA1 antibody. The data are representative of three independent experiments.

### Nuclear HBc aggregates are sensitive to proteasome inhibition and PML nuclear bodies accumulate ubiquitinylated proteins

PML nuclear bodies have been described as protective sites against nuclear proteotoxic stress by directly coordinating the proteasome-mediated degradation of aggregated proteins (Guo et al., 2014). We therefore tested whether PML nuclear bodies are associated with HBc degradation. HepG2-HBc-HA cells were treated with CAM-A for 16 hours to induce HBc nuclear aggregates and then MG132 (a proteasome inhibitor) or bafilomycin (an autophagy inhibitor) was added for 8 hours, after which the levels of HBc were measured using flow cytometry. Our results showed that only proteasome inhibition by MG132 increased the level of HBc (Figure 5A). Therefore, these findings suggest that the ubiquitin-proteasome system is the primary pathway responsible for the degradation of aggregated nuclear HBc. We also performed LC3 (marker of autophagosome) staining in HepG2-NTCP cells expressing HBc; however, no LC3 signal co-localised markedly with nuclear HBc aggregates post CAM-A treatment (supplementary figure 4). Thus, we conclude that HBc nuclear aggregates are not degraded by autophagy. PML nuclear bodies may also accumulate ubiquitinylated proteins during proteotoxic stress (Villagra et al., 2006). Next, we tested whether enlarged PML nuclear bodies and HBc aggregates colocalise with ubiquitine signal following CAM-A treatment, using an antibody that recognises both mono- and polyubiquitinated proteins. Interestingly, we observed that CAM-A treatment leads to an accumulation of ubiquitinated proteins in nuclear structures that colocalize both with PML bodies and HBc (Figure 5B), further confirming that HBc aggregates are likely to be degraded by proteasomal-ubiquitin system.

**Figure 5.**
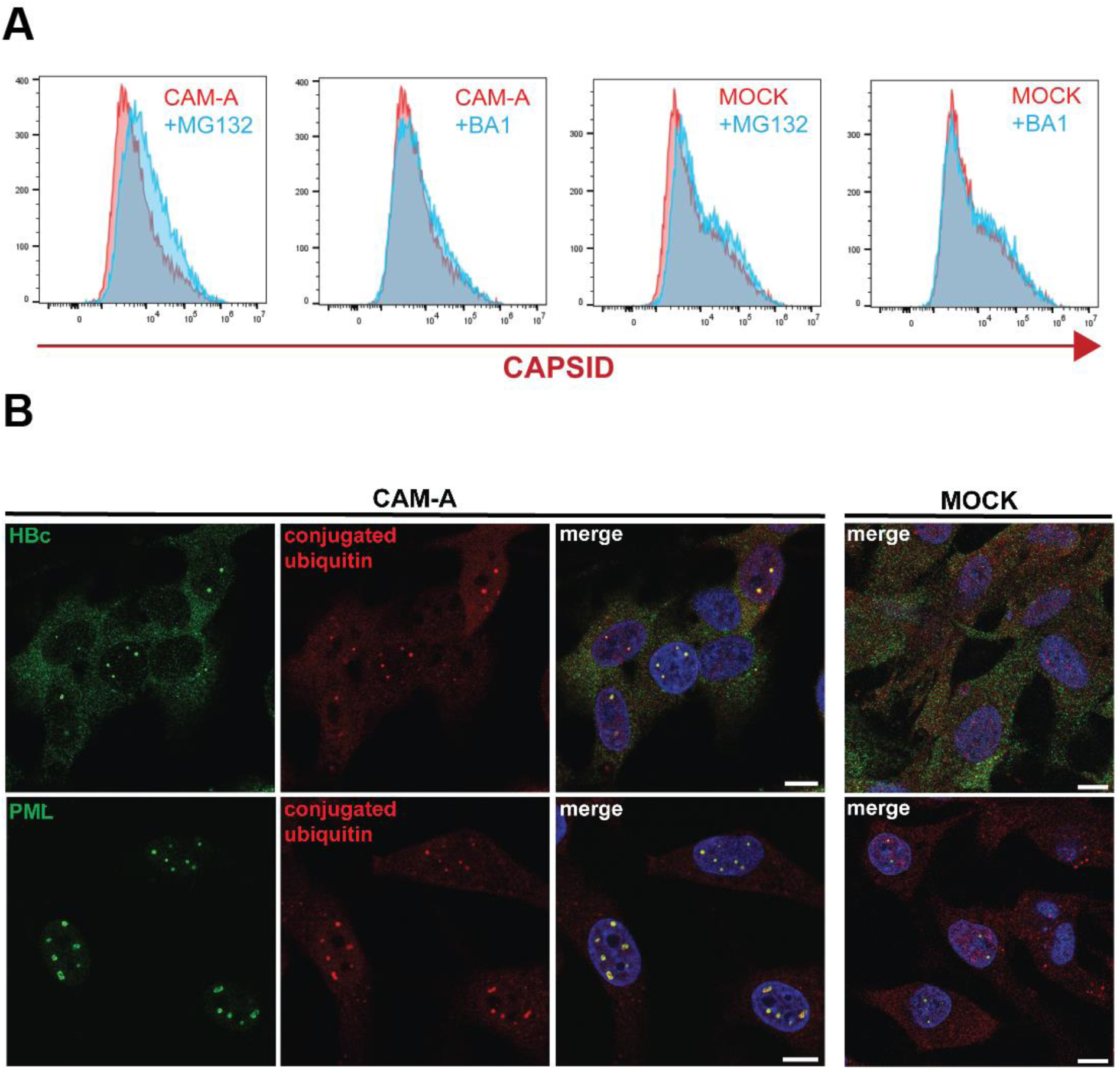
PML nuclear bodies orchestrate ubiquitin proteasome degradation of CAM-A-induced HBc aggregates. (A) HepG2-NTCP expressing HBc-HA were treated with CAM-A (RG7907; 1 µM) for 16 hours to induce nuclear HBc aggregates and then treated with MG132 or Bafilomycin A1 for 8 hours. The level of aggregated HBc was measured by flow cytometry using a capsid antibody. The data are representative of three independent experiments. (B) HepG2-NTCP expressing HBc-HA were treated with CAM-A (RG7907; 1 µM) or MOCK for 24 hours. PML (green) or HBc (green) and ubiquitinylated proteins (red) were stained by specific antibodies and analysed by confocal microscopy. The cell nucleus was stained by DAPI (blue). Bars = 10 μm.

### PML nuclear bodies orchestrate HBc degradation via the recruitment of the SUMO-targeted ubiquitin ligase RNF4

The function and composition of PML nuclear bodies are regulated by interactions with proteins via sumoylation (Shen et al., 2006). We therefore tested whether sumoylation also plays a role in CAM-A-mediated HBc aggregation. We examined the formation of HBc aggregates induced by CAM-A in cells treated with TAK-981, a sumoylation inhibitor, using confocal microscopy. Our data indicate that inhibition of sumoylation does not affect CAM-A-mediated nuclear HBc aggregation or its association with PML nuclear bodies (Figure 6A). However, the size of HBc aggregates increased markedly. PML nuclear bodies have been shown to coordinate the degradation of misfolded proteins by recruiting the E3 ubiquitine ligase RNF4, which ubiquitinates proteins for degradation (Guo et al., 2014). RNF4 belongs to a family of SUMO-targeted ubiquitin ligases (STUbLs), which mediate the ubiquitination of sumoylated proteins (Xu et al., 2014). First, we analysed whether RNF4 localizes to PML nuclear bodies following CAM-A treatment using confocal microscopy. We confirmed that RNF4 is a component of enlarged PML nuclear bodies (Figure 6B). We therefore silenced RNF4 expression protein using siRNA and tested the effect of CAM-A in HBc-expressing cells. Flow cytometry revealed that the level of HBc increased in cells with silenced RNF4 compared to the control (Fig. 6B). Our results suggest that PML nuclear bodies orchestrate the degradation of aggregated HBc via RNF4 recruitment.

**Figure 6.**
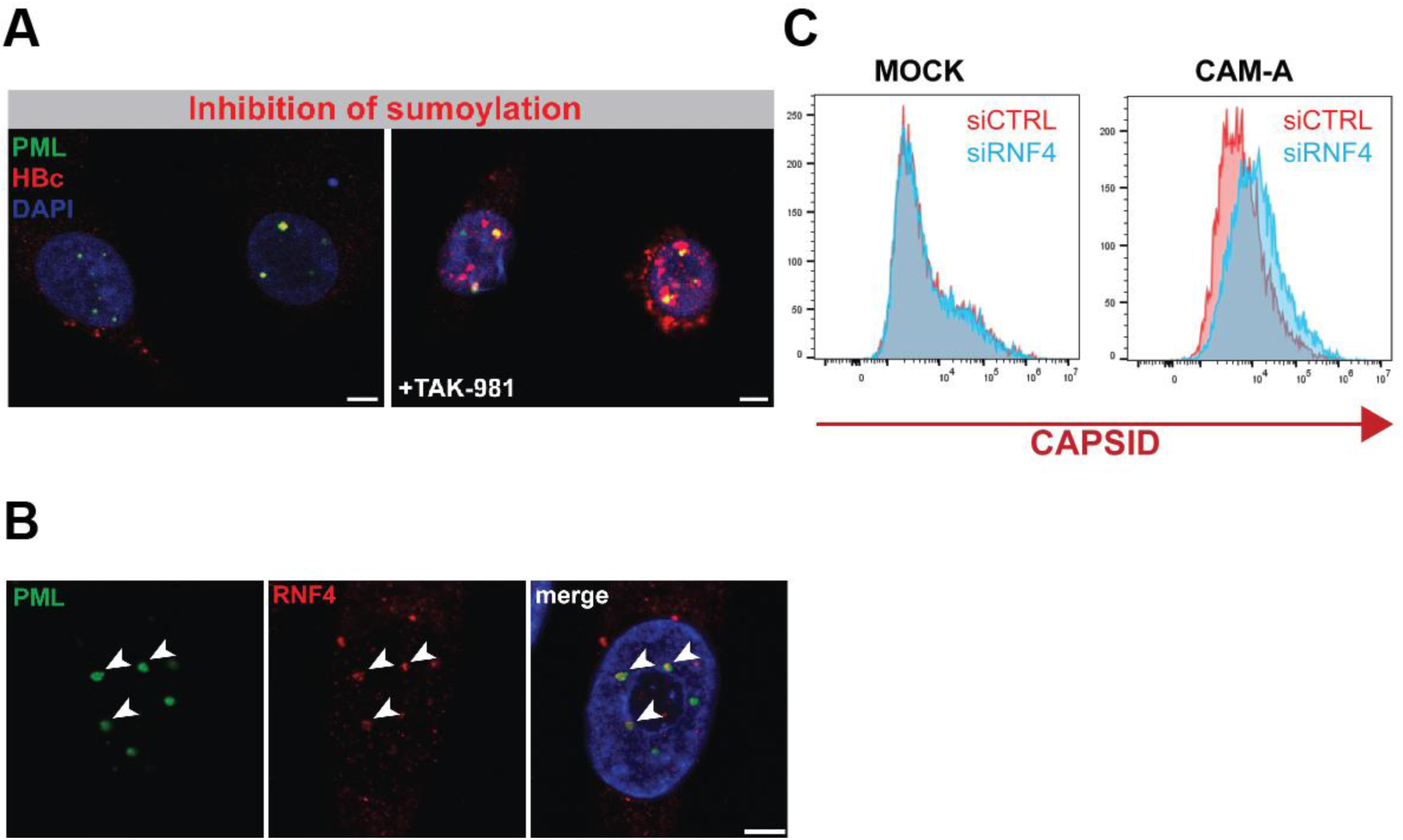
PML nuclear bodies orchestrate ubiquitin proteasome degradation of CAM-A-induced HBc aggregates. HepG2-NTCP expressing HBc-HA were treated with CAM-A (RG7907; 1 µM) for 24 hours to induce nuclear HBc aggregates and then treated with TAK-981 (5 µM) for 24 hours. PML (green) and HBc (green) were stained with specific antibodies and analysed by confocal microscopy. Bars = 5 μm (B) The localization of RNF4 (red) and PML (green) in CAM-A-treated (RG7907; 1 µM) HepG2-NTCP cells expressing HBc-HA was analysed by confocal microscopy. White arrows show areas of PML and RNF4 colocalizations. Bar = 5 μm. The nucleus was stained by DAPI. (C) Levels of HBc in CAM-A-treated (RG7907; 1 µM) HepG2-NTCP expressing HBc-HA (RG7907; 1 µM) transfected with RNF4 or control siRNA were measured by flow cytometry. The data are representative of two independent experiments.

## Discussion

Despite the previously described association between CAM-A-induced HBc aggregates and PML nuclear bodies, the precise mechanism has not been defined (Vanrusselt et al., 2023; Huber et al., 2018). We demonstrated previously that CAM-A induces the apoptosis of HBc-expressing hepatocytes via HBc nuclear aggregation and ANXA1 upregulation (Taverniti et al., 2024). In this study, we further investigated the role of PML nuclear bodies in the apoptotic elimination of HBc-containing hepatocytes by CAM-A. PML nuclear bodies play important roles in several cellular processes, including the proteasomal degradation of protein aggregates and apoptosis (Dorosz et al., 2025; Gärtner and Muller, 2014; Wang et al., 1998). Thus, we investigated whether the association of HBc aggregates with PML nuclear impacts ANXA1-driven apoptosis.

First, we observed that CAM-A promotes the accumulation of nuclear HBc aggregates within PML nuclear bodies in HBc-expressing cells. Nuclear HBc aggregates preferentially localise to enlarged PML bodies, suggesting that CAM-A treatment leads to the structural reorganization of these bodies. Large, ring-shaped PML bodies that trap viral capsids have previously been described in varicella-zoster virus and human cytomegalovirus infections as part of the antiviral mechanism. These structures are called giant PML bodies or PML cages (Reichelt et al., 2011; Scherer et al., 2022). Moreover, it was shown that interferon and DNA damage signalling induce the formation of giant PML bodies (Scherer et al., 2022). We have shown that also aggregated HBc is trapped in enlarged PML resembling PML cages.

We clearly demonstrated that PML nuclear bodies protect against CAM-A-induced apoptosis and are not the drivers of ANXA-1-mediated apoptosis. Our results suggest that HBc aggregates are targeted by PML to proteasomal degradation. Several reports demonstrated that HBc is degraded by the proteasome ubiquitin system (Braun et al., 2007; Nakaya et al., 2023; Qian et al., 2012). We also observed that, while cytoplasmic HBc aggregates were rapidly cleared, the nuclear aggregates persisted during long-term CAM-A treatment. One possible explanation is that, while HBc aggregates are effectively cleared from the cytoplasm by autophagy, as previously described (Lin et al., 2022), they may persist in the nucleus, where the degradation is primarily mediated by the proteasome.

Nuclear aggregates may affect integrity of the nucleus by inducing DNA damage and affecting nuclear transport (Gasset-Rosa et al., 2017; Korsten et al., 2024). Thus, PML nuclear bodies possibly play a dual role in maintaining the integrity of the nucleus during proteotoxic stress by directly orchestrating proteasomal degradation, and also by storing insoluble aggregates during long term proteotoxic stress when the proteasomal system in the nucleus is overwhelmed (Mediani et al., 2019; Wagner et al., 2025). CAM-A-induced HBc aggregates are highly insoluble and require harsh cell lysis conditions for extraction (Vanrusselt et al., 2023). Our confocal microscopy analysis revealed that the CAM-A-aggregated HBc is localised in the centre of the enlarged PML body. It has been shown that the centre of the PML bodies does not contain any DNA (Ryabchenko et al., 2023) and thus represents a potential storage centre where detrimental aggregates are physically separated from the DNA. Mediani et al., also proposed a model where long-lasting proteotoxic stress converts PML nuclear bodies into an amyloid-like solid state that immobilizes ubiquitin and proteasomes further confirming the storage capacity of PML nuclear bodies (Mediani et al., 2019). Further studies on PML nuclear body dynamics is required to determine whether enlarged PML bodies serve as a temporary location for the degradation of HBc aggregates or as a long-term storage site for aggregated HBc.

Lastly, our results indicate that PML-mediated degradation of nuclear HBc aggregates depends on sumoylation-dependent targeting of PML-associated proteins. E3 ubiquitin ligase RNF4 contains four tandem SUMO interaction motifs (SIMs) and has a high affinity for sumoylated proteins. After recognising sumoylated proteins, RNF4 binds to them via a SUMO-SIM interaction and ubiquitinates them for degradation (Gärtner and Muller, 2014). Therefore, the interplay between sumoylation and ubiquitylation is an important factor in managing proteotoxic stress in the nucleus (Guo et al., 2014). Furthermore, RNF4 binds and ubiquitinates PML, thereby limiting the number of PML nuclear bodies. Therefore, the entire PML body containing aggregated HBc may be targeted for degradation by RNF4. Alternatively, other E3 ubiquitin ligases associated with PML nuclear bodies, such as TOPORS or RNF111 (Van Damme et al., 2010), may also ubiquitinate the core for proteasomal degradation.

Collectively, we propose a model where PML nuclear bodies orchestrate both the degradation and storage of CAM-A-induced HBc aggregates and protects cells against CAM-A induced apoptosis.

## Materials and methods

### Cell lines

HepG2-NTCP cells were maintained in Dulbecco’s modified Eagle’s medium (DMEM, Gibco, Thermo Fisher Scientific, USA) supplemented with 10% fetal bovine serum (FBS, Dutscher, France), 1x non-essential amino acids (NEAA, Gibco, Thermo Fisher Scientific, USA), 50 μg/ml gentamicin (Gibco, Thermo Fisher Scientific, USA), and 250 μg/ml G418 (Invivogen, Thermo Fisher Scientific, USA). HBc-HA WT and HBc-HA T33N mutant-overexpressing HepG2-NTCP cells were prepared and cultivated as described previously (Taverniti et al., 2024).

### HBV production and infection

Infectious HBV particles (ayw) were prepared using HepAD38 cell and viral inoculum was quantified as previously described (Taverniti et al., 2024). HepG2-NTCP cells were infected with HBV at a multiplicity of infection (MOI) of 1000 viral genome equivalents per cell (vge/cell) in presence of 4% PEG.

### Reagents and antibodies

Capsid assembly modulators CAM-A RG7907, ALG-005398, ALG-005863 were provided by Aligos Therapeutics (Leuven, Belgium). Compound B (Aligos Therapeutics) served as a reference CAM-E and was used as previously described (Taverniti et al., 2024). All CAMs were reconstituted in DMSO (Sigma-Aldrich, Merck, Germany). TAK-981, Bafilomycin A1 and MG-132 were purchased from MedChemExpress (New Jersey, USA). Mouse monoclonal antibodies targeting human PML were purchased from Santa Cruz (Dallas, TX, USA) and rabbit monoclonal antibody targeting PML was purchased from Cell Signaling Technology (Danvers, MA, USA). Rabbit monoclonal antibody against LC3B was purchased from Cell Signaling Technology (Danvers, MA, USA). Mouse monoclonal antibodies specific against HBV capsids (Hyb-3120) were purchased from the Institute of Immunology Co. (Japan), LTD. Anti-HA tag antibody was obtained from Abcam (Cambridge, UK). Mouse antibody targeting ubiquitinylated proteins was obtained from Merck (Darmstadt, Germany). Rabbit monoclonal antibody detecting Annexin A1/ANXA1 was obtained from Abcam (Cambridge, UK). Rabbit polyclonal antibody targeting RNF4 was purchased from Thermo Fisher Scientific (Waltham, USA).

### Cell viability and apoptosis assays

The rate of apoptosis was performed as described previously (Taverniti et al., 2024). Briefly, Cells were seeded in 96-well plate and treated with various CAM-A for 3 days. Then, cells were incubated with the Cell Event Caspase-3/7 Green Detection Reagent (Invitrogen, Thermo Fisher Scientific, USA, C10723) following the manufacturer’s instructions. Quantification was made using a Celigo Image Cytometer.

### siRNA and cell transfection

The cells were reverse transfected with specific siRNA using RNAiMAX (Thermo Fisher Scientific), according to the manufacturer’s protocol. The transfection reagent was replaced with fresh medium six hours after transfection. SMARTpoo siRNA against PML and RNF4, including non-targeting control siRNA, was purchased from Dharmacon. PML and RNF4 expression was analysed 48 hours post-transfection using confocal microscopy and RT-qPCR.

### Analysis of ANXA1 and aggregated HBc by Flow Cytometry

Cells were trypsinized and fixed with 4% formaldehyde in PBS for 15 min. Cells were then permeabilized with 0,5% Triton for 5 minutes and washed 3× with PBS. Cells were incubated with primary antibodies in 1% BSA for 1 hour at room temperature. As secondary antibody served donkey anti-rabbit conjugated to Alexa Fluor 488 (Thermo Fisher Scientific) and goat anti-mouse conjugated to Alexa Fluor 647 (Thermo Fisher Scientific). Samples were analyzed using a CytoFLEX Flow Cytometer (Beckman Coulter, Brea, California, United States), and data were processed using FLOWJO software (Treestar, San Carlos, CA, USA).

### Immunofluorescence assay (IFA) and confocal microscopy

Cells were washed with PBS and fixed with 4% formaldehyde in PBS for 15 min. Cells were then permeabilized with 0.5% Triton X-100 in PBS for 5 min, washed 3× with PBS and blocked with 1% BSA for 1 h at room temperature. Specific primary antibodies were diluted in 1% BSA and were added for 1 h at room temperature. Donkey anti-rabbit conjugated to Alexa Fluor 488 and goat anti-mouse conjugated to Alexa Fluor 647 were used as secondary antibodies. Cell nuclei were stained with DAPI. Cells were then mounted on droplets of SlowFade Diamond Antifade Mountant (Thermo Fisher Scientific). Images were captured using a Zeiss LSM 800 AiryScan (Carl Zeiss). Fiji and Zeiss Zen softwares were used for data evaluation. Fiji was used for the measurement of HBc aggregates.

### Quantification of gene expression by RT-qPCR

Total RNA was isolated using the ReliaPrep RNA Miniprep Systems (Promega, USA) and then reverse-transcribed into complementary DNA (cDNA) using the Maxima First Strand cDNA Synthesis Kit (Thermo Fisher Scientific, USA) according to the manufacturer’s instructions. Gene expression was then quantified by quantitative PCR using an Applied Biosystems instrument. Primers for PML and GAPDH mRNA quantification were obtained from Sigma-Aldrich. Sequence of PML primers: forward: 5′- CACCCGCAAGACCAACAACA-3′ and reverse 5′- ATCCTCGGCAGTAGATGCTGG-3′. GAPDH primers were used as described previously (Verrier et al., 2023). All values were normalized to GAPDH expression. Relative expression levels were calculated using the 2^−ΔΔCT^ method.

### Data analysis and statistics

Quantitative variables are expressed as the mean ± standard error of the mean (SEM). The number of independent experiments per assay is indicated in the figure legends. Representative images and results were selected from confocal microscopy and flow cytometry and presented. Data were analyzed with GraphPad Prism 6 (GraphPad Software). A p-value of ≤0.05 was considered significant.

## Authors’ contributions

Study concept and design: VJ and ERV. Study supervision: ERV. Acquisition of data: VJ with the support of LM-H, VT, and CS. Analysis and interpretation of data: VJ, JW, BL, JL, IH, TFB, and ERV. Material support and discussion: HV and YD. Funding acquisition: ERV, VJ, VT, TFB, and YD. Drafting of the manuscript: VJ and ERV. All the authors approved the manuscript.

## Financial support

This work of the Interdisciplinary Thematic Institute IMCBio, as part of the ITI 2021-2028 program of the University of Strasbourg, CNRS and Inserm, was supported by IdEx Unistra (ANR-10-IDEX-0002), and by SFRI-STRAT’US project (ANR-20-SFRI-0012) and EUR IMCBio (ANR-17-EURE-0023) under the framework of the French Investments for the Future Program. TFB and ERV received funding from Aligos Belgium BV as part of the VLAIO project CoHeBA (HBC.2020.2454). VT and ERV acknowledge fundings from ANRS - Maladies infectieuses émergentes - (ANRS MIE, grant number ANRS0543). TFB acknowledges funding from the European Union (EUERC-AdG-2014-HEPCIR #671231) and ARC Foundation TheraHCC2.0 (IHU201901299). VJ acknowledges funding from Czech science foundation Grant No. 24-11554O (Identification of novel antiviral targets in the nucleus of hepatitis B virus infected cells).

## Data availability statement

The original data from this study are available through the corresponding author upon reasonable request.

## Abbreviations

CAMs: Capsid assembly modulators
cccDNA: covalently closed circular DNA (cccDNA)
DMSO: dimethyl sulfoxide
HBc: HBV core antigen
NEAA: non-essential amino acids
NUCs: nucleos(t)ide analogues
pgRNA: pre-genomic RNA
PML: Promyelocytic leukemia protein
RNF4: RING finger protein 4
SIM: SUMO interacting motif
SUMO: Small ubiquitin-related modifier
Vge: viral genome equivalents

## Acknowledgments

We thank CRBS Imaging Platform “PIC-STRA” (University of Strasbourg) and our colleagues Ms Sarah Durand (U1110) and Ms Marine Oudot (U1110) for excellent technical support. We thank our colleague Ms Anne Zeter (U1110) for excellent administrative support.

## Conflicts of interest

TFB and ERV received funding from Aligos Belgium BV as part of the VLAIO project CoHeBA (HBC.2020.2454). YD, and HV are employees of Aligos and may own stock.

## References

Berke, J.M., Dehertogh, P., Vergauwen, K., Van Damme, E., Mostmans, W., Vandyck, K., Pauwels, F., 2017. Capsid Assembly Modulators Have a Dual Mechanism of Action in Primary Human Hepatocytes Infected with Hepatitis B Virus. Antimicrob Agents Chemother 61, e00560–17. 10.1128/AAC.00560-17

Berke, J.M., Tan, Y., Sauviller, S., Wu, D.-T., Zhang, K., Conceição-Neto, N., Blázquez Moreno, A., Kong, D., Kukolj, G., Li, C., Zhu, R., Nájera, I., Pauwels, F., 2024. Class A capsid assembly modulator apoptotic elimination of hepatocytes with high HBV core antigen level in vivo is dependent on de novo core protein translation. J Virol 98, e0150223. 10.1128/jvi.01502-23

Braun, S., Zajakina, A., Aleksejeva, J., Sharipo, A., Bruvere, R., Ose, V., Pumpens, P., Garoff, H., Meisel, H., Kozlovska, T., 2007. Proteasomal degradation of core protein variants from chronic hepatitis B patients. J Med Virol 79, 1312–1321. 10.1002/jmv.20939

Dorosz, K., Majewska, L., Kijowski, J., 2025. Structure and Function of PML Nuclear Bodies: A Brief Overview of Key Cellular Roles. Biomolecules 15. 10.3390/biom15091291

Gärtner, A., Muller, S., 2014. PML, SUMO, and RNF4: Guardians of Nuclear Protein Quality. Molecular Cell 55, 1–3. 10.1016/j.molcel.2014.06.022

Gasset-Rosa, F., Chillon-Marinas, C., Goginashvili, A., Atwal, R.S., Artates, J.W., Tabet, R., Wheeler, V.C., Bang, A.G., Cleveland, D.W., Lagier-Tourenne, C., 2017. Polyglutamine-Expanded Huntingtin Exacerbates Age-Related Disruption of Nuclear Integrity and Nucleocytoplasmic Transport. Neuron 94, 48–57.e4. 10.1016/j.neuron.2017.03.027

Glebe, D., Goldmann, N., Lauber, C., Seitz, S., 2021. HBV evolution and genetic variability: Impact on prevention, treatment and development of antivirals. Antiviral Res 186, 104973. 10.1016/j.antiviral.2020.104973

Guo, L., Giasson, B.I., Glavis-Bloom, A., Brewer, M.D., Shorter, J., Gitler, A.D., Yang, X., 2014. A cellular system that degrades misfolded proteins and protects against neurodegeneration. Mol Cell 55, 15–30. 10.1016/j.molcel.2014.04.030

Hsieh, C.-L., Chang, L.-Y., Chen, P.-J., Yeh, S.-H., 2025. HBV polymerase recruits the phosphatase PP1 to dephosphorylate HBc-Ser170 to complete encapsidation. PLOS Pathogens 21, e1012905. 10.1371/journal.ppat.1012905

Huber, A.D., Wolf, J.J., Liu, D., Gres, A.T., Tang, J., Boschert, K.N., Puray-Chavez, M.N., Pineda, D.L., Laughlin, T.G., Coonrod, E.M., Yang, Q., Ji, J., Kirby, K.A., Wang, Z., Sarafianos, S.G., 2018. The Heteroaryldihydropyrimidine Bay 38-7690 Induces Hepatitis B Virus Core Protein Aggregates Associated with Promyelocytic Leukemia Nuclear Bodies in Infected Cells. mSphere 3, 10.1128/mspheredirect.00131-18. 10.1128/mspheredirect.00131-18

Kennedy, P.T., Allweiss, L., Bertoletti, A., Cornberg, M., Gehring, A.J., Guidotti, L.G., Kerth, H.A., Lemoine, M., Levrero, M., Lim, S.G., Tavis, J.E., Testoni, B., Tu, T., 2025. Scientific and medical evidence informing expansion of hepatitis B treatment guidelines. The Lancet Gastroenterology & Hepatology 10, 941–951. 10.1016/S2468-1253(25)00053-6

Korsten, G., Osinga, M., Pelle, R.A., Serweta, A.K., Hoogenberg, B., Kampinga, H.H., Kapitein, L.C., 2024. Nuclear poly-glutamine aggregates rupture the nuclear envelope and hinder its repair. J Cell Biol 223, e202307142. 10.1083/jcb.202307142

Kum, D.B., Vanrusselt, H., Acosta Sanchez, A., Taverniti, V., Verrier, E.R., Baumert, T.F., Liu, C., Deval, J., Corthout, N., Munck, S., Beigelman, L., Blatt, L.M., Symons, J.A., Raboisson, P., Jekle, A., Vendeville, S., Debing, Y., 2023. Class A capsid assembly modulator RG7907 clears HBV-infected hepatocytes through core-dependent hepatocyte death and proliferation. Hepatology 78, 1252–1265. 10.1097/HEP.0000000000000428

Lin, J., Yin, L., Xu, X.-Z., Sun, H.-C., Huang, Z.-H., Ni, X.-Y., Chen, Y., Lin, X., 2022. Bay41-4109-induced aberrant polymers of hepatitis b capsid proteins are removed via STUB1-promoted p62-mediated macroautophagy. PLOS Pathogens 18, e1010204. 10.1371/journal.ppat.1010204

Lott, L., Beames, B., Notvall, L., Lanford, R.E., 2000. Interaction between Hepatitis B Virus Core Protein and Reverse Transcriptase. J Virol 74, 11479–11489. 10.1128/jvi.74.24.11479-11489.2000

Lubyova, B., Tikalova, E., Janovec, V., Ryabchenko, B., Krulova, K., Kropacek, V., Huerfano, S., Hirsch, I., Weber, J., 2025. UBE2O, a host ubiquitin-conjugating enzyme, is a key regulator of hepatitis B virus maturation and egress. J Biol Chem 301, 110750. 10.1016/j.jbc.2025.110750

Luo, J., Xi, J., Gao, L., Hu, J., 2020. Role of Hepatitis B virus capsid phosphorylation in nucleocapsid disassembly and covalently closed circular DNA formation. PLoS Pathog 16, e1008459. 10.1371/journal.ppat.1008459

Mediani, L., Guillén-Boixet, J., Alberti, S., Carra, S., 2019. Nucleoli and Promyelocytic Leukemia Protein (PML) bodies are phase separated nuclear protein quality control compartments for misfolded proteins. Mol Cell Oncol 6, e1415624. 10.1080/23723556.2019.1652519

Nakaya, Y., Nishizawa, T., Nishitsuji, H., Morita, H., Yamagata, T., Onomura, D., Murata, K., 2023. TRIM26 positively affects hepatitis B virus replication by inhibiting proteasome-dependent degradation of viral core protein. Sci Rep 13, 13584. 10.1038/s41598-023-40688-3

Niklasch, M., Zimmermann, P., Nassal, M., 2021. The Hepatitis B Virus Nucleocapsid-Dynamic Compartment for Infectious Virus Production and New Antiviral Target. Biomedicines 9, 1577. 10.3390/biomedicines9111577

Qian, G., Jin, F., Chang, L., Yang, Y., Peng, H., Duan, C., 2012. NIRF, a novel ubiquitin ligase, interacts with hepatitis B virus core protein and promotes its degradation. Biotechnol Lett 34, 29–36. 10.1007/s10529-011-0751-0

Reichelt, M., Wang, L., Sommer, M., Perrino, J., Nour, A.M., Sen, N., Baiker, A., Zerboni, L., Arvin, A.M., 2011. Entrapment of Viral Capsids in Nuclear PML Cages Is an Intrinsic Antiviral Host Defense against Varicella-Zoster Virus. PLoS Pathog 7, e1001266. 10.1371/journal.ppat.1001266

Ryabchenko, B., Šroller, V., Horníková, L., Lovtsov, A., Forstová, J., Huérfano, S., 2023. The interactions between PML nuclear bodies and small and medium size DNA viruses. Virol J 20, 82. 10.1186/s12985-023-02049-4

Scherer, M., Read, C., Neusser, G., Kranz, C., Kuderna, A.K., Müller, R., Full, F., Wörz, S., Reichel, A., Schilling, E.-M., Walther, P., Stamminger, T., 2022. Dual signaling via interferon and DNA damage response elicits entrapment by giant PML nuclear bodies. eLife 11, e73006. 10.7554/eLife.73006

Shen, T.H., Lin, H.-K., Scaglioni, P.P., Yung, T.M., Pandolfi, P.P., 2006. The Mechanisms of PML-Nuclear Body Formation. Mol Cell 24, 331–339. 10.1016/j.molcel.2006.09.013

Taverniti, V., Ligat, G., Debing, Y., Kum, D.B., Baumert, T.F., Verrier, E.R., 2022. Capsid Assembly Modulators as Antiviral Agents against HBV: Molecular Mechanisms and Clinical Perspectives. J Clin Med 11, 1349. 10.3390/jcm11051349

Taverniti, V., Meiss-Heydmann, L., Gadenne, C., Vanrusselt, H., Kum, D.B., Giannone, F., Pessaux, P., Schuster, C., Baumert, T.F., Debing, Y., Verrier, E.R., 2024. CAM-A-dependent HBV core aggregation induces apoptosis through ANXA1. JHEP Rep 6, 101134. 10.1016/j.jhepr.2024.101134

Van Damme, E., Laukens, K., Dang, T.H., Van Ostade, X., 2010. A manually curated network of the PML nuclear body interactome reveals an important role for PML-NBs in SUMOylation dynamics. Int J Biol Sci 6, 51–67. 10.7150/ijbs.6.51

Vanrusselt, H., Kum, D.B., Taverniti, V., Liu, C., Acosta Sanchez, A., Corthout, N., Munck, S., Baumert, T.F., Beigelman, L., Blatt, L.M., Symons, J.A., Deval, J., Raboisson, P., Verrier, E.R., Jekle, A., Vendeville, S., Debing, Y., n.d. Novel non-HAP class A HBV capsid assembly modulators have distinct in vitro and in vivo profiles. J Virol 97, e00722–23. 10.1128/jvi.00722-23

Verrier, E.R., Ligat, G., Heydmann, L., Doernbrack, K., Miller, J., Maglott-Roth, A., Jühling, F., Saghire, H.E., Heuschkel, M.J., Fujiwara, N., Hsieh, S.-Y., Hoshida, Y., Root, D.E., Felli, E., Pessaux, P., Mukherji, A., Mailly, L., Schuster, C., Brino, L., Nassal, M., Baumert, T.F., 2023. Cell-based cccDNA reporter assay combined with functional genomics identifies YBX1 as HBV cccDNA host factor and antiviral candidate target. 10.1136/gutjnl-2020-323665

Villagra, N.T., Navascues, J., Casafont, I., Val-Bernal, J.F., Lafarga, M., Berciano, M.T., 2006. The PML-nuclear inclusion of human supraoptic neurons: a new compartment with SUMO-1-and ubiquitin–proteasome-associated domains. Neurobiology of Disease 21, 181–193. 10.1016/j.nbd.2005.07.003

Wagner, K., Keiten-Schmitz, J., Adhikari, B., Patra, U., Husnjak, K., McNicoll, F., Dormann, D., Müller-McNicoll, M., Tascher, G., Wolf, E., Müller, S., 2025. Induced proximity to PML protects TDP-43 from aggregation via SUMO–ubiquitin networks. Nat Chem Biol 21, 1408–1419. 10.1038/s41589-025-01886-4

Wang, Z.G., Ruggero, D., Ronchetti, S., Zhong, S., Gaboli, M., Rivi, R., Pandolfi, P.P., 1998. PML is essential for multiple apoptotic pathways. Nat Genet 20, 266–272. 10.1038/3073

Xu, Y., Plechanovová, A., Simpson, P., Marchant, J., Leidecker, O., Kraatz, S., Hay, R.T., Matthews, S.J., 2014. Structural insight into SUMO chain recognition and manipulation by the ubiquitin ligase RNF4. Nat Commun 5, 4217. 10.1038/ncomms5217

Zhao, Q., Hu, Z., Cheng, J., Wu, S., Luo, Y., Chang, J., Hu, J., Guo, J.-T., 2018. Hepatitis B Virus Core Protein Dephosphorylation Occurs during Pregenomic RNA Encapsidation. J Virol 92, e02139–17. 10.1128/JVI.02139-17

