## Supplementary figure for "PML nuclear bodies orchestrate the storage and degradation of aggregated HBc in the nucleus and reduce CAM-A-induced apoptosis"

### SUPPLEMENTARY FIGURES

**Supplementary video 1. Z-stack imaging analysis of enlarged PML bodies in an individual cell.**
3D projection of PML nuclear bodies (single channel, green) stained with specific antibodies. Link to video:

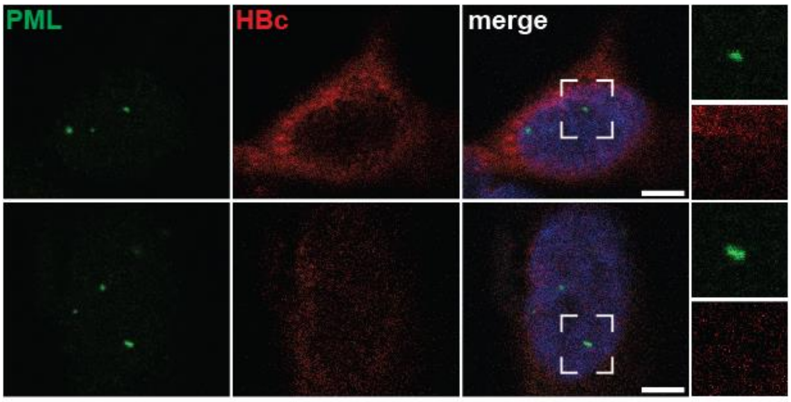


**Supplementary Figure 1.**
HepG2-HBc-HA cells were treated with CAM-E (Compound B, 1 μM; 24 hours) and stained with antibodies against PML (green) and HA-Tag (red). Nuclei were counterstained with DAPI (blue). White squares show enlarged regions. Bar = 5 μm.

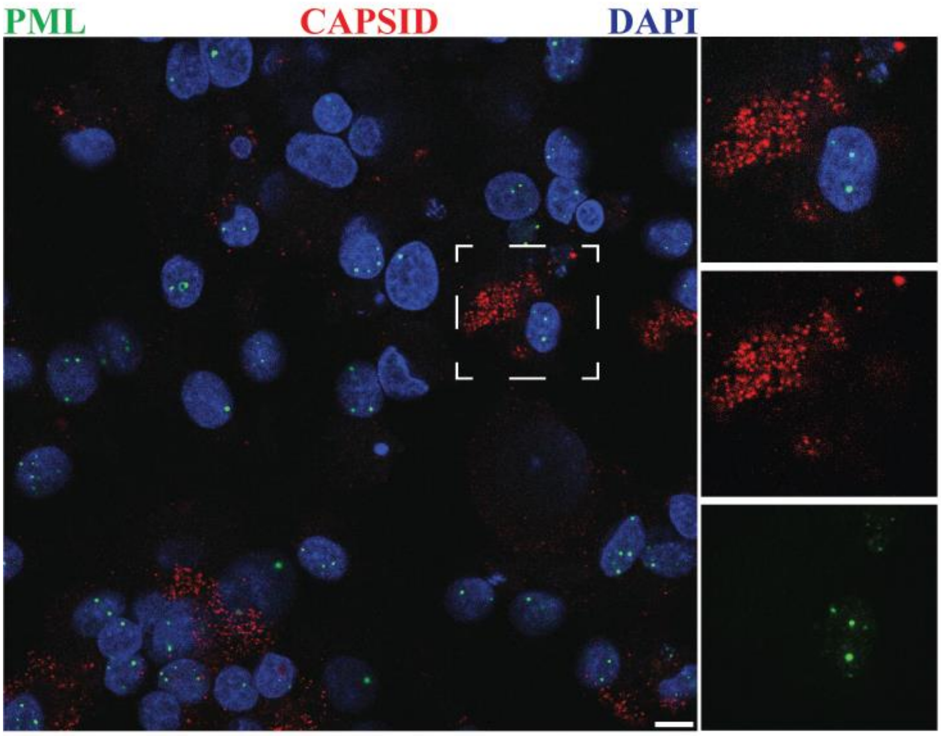


**Supplementary Figure 2.**
HepG2-NTCP cells were infected with HBV (1000GE/cell) and treated with MOCK (DMSO) for 6 days and stained with antibodies against PML (green) and HBV capsid (red). Nuclei were counterstained with DAPI (blue). White squares show enlarged regions of PML and HBV capsid signals analyzed by confocal microscopy. Bar = 10 µm.

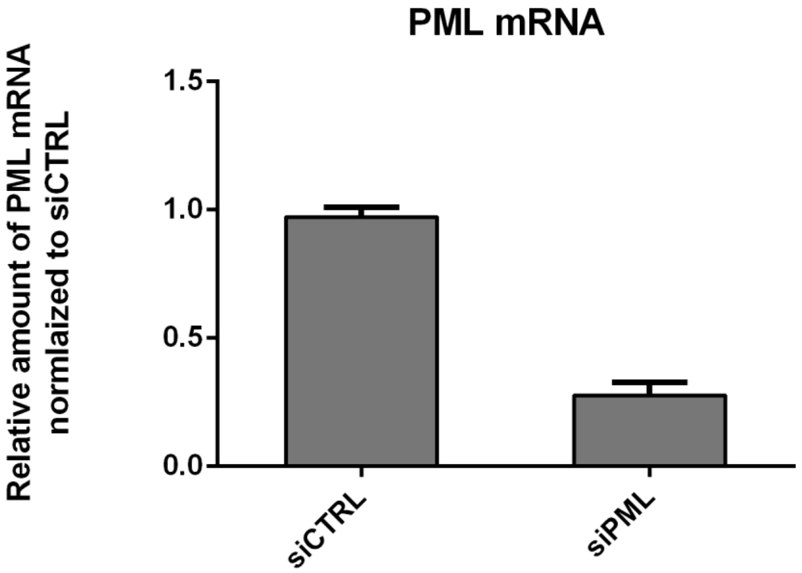


**Supplementary Figure 3**. RT-qPCR analysis of PML mRNA after transfection of siRNA targeting PML or non-targeting siRNA.

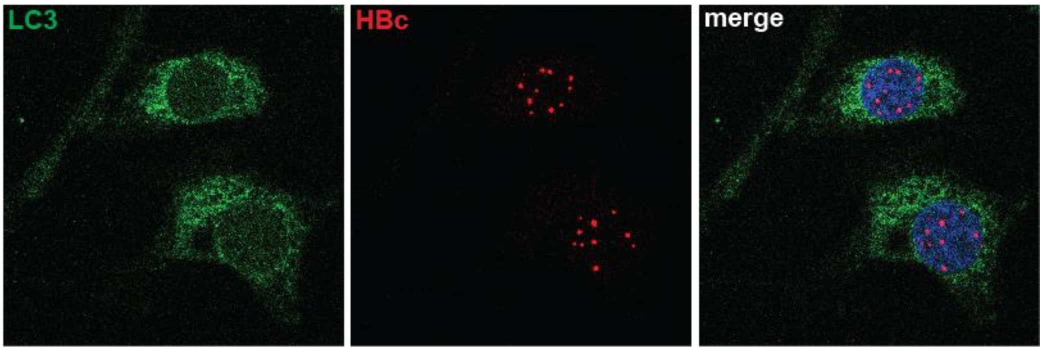


**Supplementary Figure 4**. **LC3 staining in the nucleus.** HepG2-HA-HBc cells were treated with CAM-A (ALG-005863; 1 µM; 24 hours) and stained with antibodies against LC3B (green) and HA-Tag (red). Nucleus was stained by DAPI.
